# Circumventing the synthesizability problem in generative molecular design

**DOI:** 10.64898/2026.02.18.706722

**Authors:** Jesse A. Weller, Jinsen Li, Yibei Jiang, Remo Rohs

## Abstract

Generative structure-based drug design (SBDD) models have shown great promise to accelerate our ability to discover novel drug candidates. However, these models have been criticized for producing compounds that are not very synthesizable, and therefore not practically applicable to drug design. In this work, we propose a way to circumvent the synthesizability issue by introducing a model-guided virtual screening (MGVS) pipeline which pairs SBDD models with efficient chemical similarity search methods to identify synthesizable analogs of generated compounds in existing ultra-large compound databases. Using this approach, we demonstrate that synthesizable analogs of generated compounds with equivalent or better docking scores and similar predicted binding poses can be reliably identified across a wide range of protein targets. We find that MGVS outperforms standard virtual ligand screening (VLS), consistently yielding at least a 25x improvement in screening efficiency across three different SBDD models. As drug-like chemical spaces continue to grow and standard VLS methods focused on exhaustive screening become increasingly impractical, approaches like MGVS that effectively narrow the search space will become critical for advancing drug discovery.

## INTRODUCTION

Generative deep learning methods for small-molecule drug design have captured widespread attention in both academia and industry. These approaches promise to accelerate early-stage drug discovery by identifying novel chemical structures tailored to specific biological targets, potentially unlocking areas of chemical space unexplored by traditional methods. Despite their promise, these models face a critical challenge that limits their practical utility: the synthesizability of generated compounds. This is a particular issue in the early stages of the drug discovery process, where the ability to rapidly procure and validate potential hit compounds is essential, and custom synthesis of novel chemical structures at scale is impractically slow and expensive.

For this reason, traditional virtual ligand screening (VLS) approaches have focused on screening commercially available libraries of compounds with known synthesis routes^1–4^. Historically, such libraries have been limited to only a few million compounds, whereas drug-like chemical space is on the order of 10^60^ compounds^5^. This limitation has been partially addressed with the development of virtual chemical spaces^1^—enumerated or reaction-defined collections of make-on-demand compounds that can be synthesized reliably at scale. Public databases now include billions^2^ to trillions^6^ of compounds, constructed by applying known reactions to commercially available building blocks.

While these advances have dramatically expanded the accessible chemical search space, they have also introduced the daunting computational challenge of screening chemical spaces of this scale. Parallel advancements in VLS methods such as synthon-based screening^7,8^ have lowered the computational overhead by systematically docking chemical building blocks instead of every reachable compound, thus helping to mitigate the combinatorial explosion. As virtual spaces continue to grow, however, such knowledge-free approaches that rely on exhaustively screening the chemical space will become increasingly impractical. In recognition of this challenge, recent efforts have turned toward advanced sampling strategies—ranging from traditional heuristics to modern AI-based methods^1,9,10^ to narrow the search space by screening a small fraction of the chemical space and using the most promising candidates to guide further search. However, these approaches remain constrained by their dependence on large-scale enumeration and iterative evaluation, which limits their efficiency and scalability as chemical spaces expand.

Generative deep learning models have recently emerged as a compelling alternative. These models can learn distributions of chemical structures and protein–ligand interactions^11–13^ from large chemical datasets, offering the potential to navigate chemical space without explicit enumeration^11,13^. When conditioned on structural information from a protein target, these models can generate compounds that fit within the protein pocket and exhibit favorable binding properties. Despite their promise, generative methods have been criticized for producing compounds that are either synthetically inaccessible or chemically implausible^14–16^. In response, there has been a focus on restricting models to explicitly learn distributions over synthesizable subspaces^17–19^, but these methods often result in making trade-offs in terms of molecular diversity and controllability. In addition, the concept of a “synthesizable” compound is not well-defined *ab initio* in terms of chemical structure or properties but instead depends on the ever-changing set of available chemical synthesis ingredients and techniques. Restricting models to such arbitrary subspaces is not straightforward and could detract from the primary goal: generating true target-specific binders. Alternatively, we hypothesize that these synthesizability issues could be circumvented by pairing generative models with established chemical similarity search techniques to identify synthesizable analogs. In this way, rather than attempting to supplant the traditional discovery pipeline by directly generating ideal, synthesizable drug candidates, generative methods may be more effectively used to steer discovery toward tractable, high-potential subspaces—within which synthesizable compounds can be identified.

In this work, we introduce a model-guided virtual screening (MGVS) pipeline that involves using structure-based generative models to sample compounds conditioned on a target pocket, selecting the compounds with strongest binding affinity, and retrieving similar compounds from existing chemical databases. We show that synthesizable compounds with strong predicted binding affinity and similar docking poses can be reliably identified using this generate-then-retrieve approach, and that the quality of the discovered candidates represents at least a 25x improvement in screening efficiency over standard VLS. These results are consistent across three state-of-the-art generative SBDD models with diverse architectures: DrugHIVE^11^ (hierarchical VAE), Pocket2Mol^12^ (autoregressive), and DiffSBDD^13^ (diffusion), exhibiting the broad applicability of this approach. In summary, this work demonstrates that: 1) existing generative SBDD models reliably identify high potential chemical subspaces and thus narrow the search for high quality candidates and 2) though not always directly synthesizable, generated compounds tend to have synthesizable analogs identifiable via similarity search. The synthesizability issue therefore does not represent a significant obstacle to their effective application to drug discovery. Improving the capacity of these models to generate high quality target-specific binders, regardless of synthesizability, could further increase the effectiveness of MGVS over traditional screening methods.

## METHODS AND DATA

An overview of our MGVS pipeline for *de novo* molecular generation followed by synthesizable analog search is shown in Figure 1. Compounds are generated using three top SBDD models with a range of architectures: DrugHIVE^11^, a density-based hierarchical variational autoencoder model; Pocket2Mol^12^, a graph-based equivariant autoregressive model; and DiffSBDD^13^, a graph-based equivariant diffusion model. For each model and target, the following five step MGVS process was used: 1) The target pocket information is provided to the generative SBDD model and used to generate 1000 compounds using default settings. 2) Compounds are then docked to the target pocket and scored using QuickVina2^20^. 3) Compounds are filtered to remove those with PAINS^21^ patterns, poor drug-like properties or irregular structures. The top-10 scoring compounds that remain are selected as queries for analog search. 4) For each of the top-10 generated query compounds, hierarchical graph edit distance (GED) similarity search is performed using SmallWorld^22^ to identify the 100 most similar compounds from existing ultra-large libraries: Enamine REAL, WuXi GalaXi, and ZINC. 5) These compounds are docked to target pocket and scored for predicted binding affinity using QuickVina2^20^. The best compounds, representing synthesizable analogs, are identified for each generated query based on docking score.

**Figure 1.**
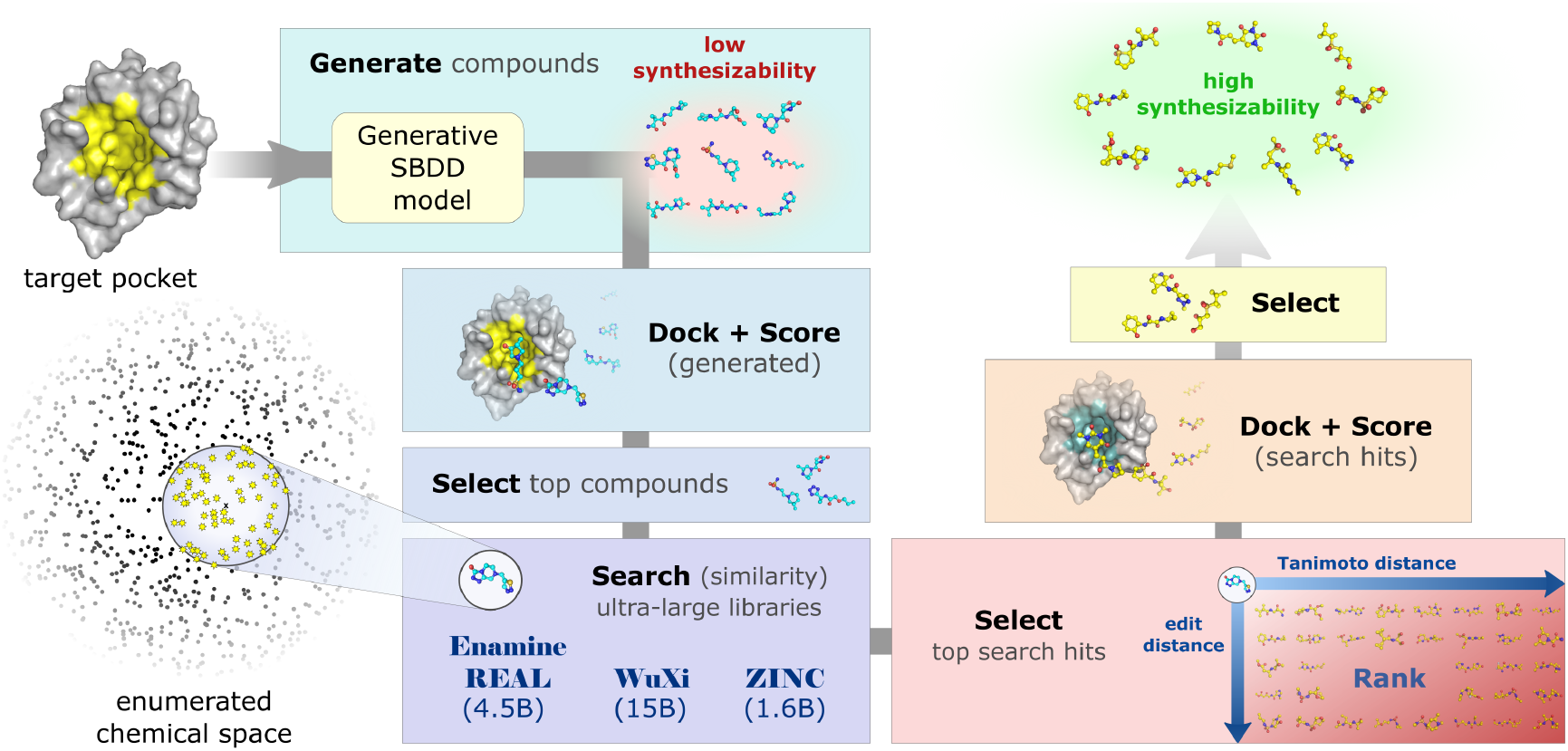
Overview of de novo generation and synthesizable analog search pipeline. 1) Target pocket information is input into the generative SBDD model and used to generate 1000 compounds. 2) Generated compounds are docked to the target pocket and scored for predicted binding affinity. 3) Compounds are filtered based on chemical structure and properties, and top-10 scoring compounds are selected for analog search. 4) For each generated query compound, efficient hierarchical graph edit distance (GED) similarity search is performed to identify the 100 most similar compounds from existing ultra-large libraries Enamine REAL, ZINC, WuXi GalaXi. 5) Top-100 most similar search-hits per query are docked to target pocket and scored for predicted binding affinity. The best synthesizable analogs for each generated query are selected based on docking.

### Compound Filtering

To avoid inaccurate docking results, we filter out molecules with irregular configurations that lead to strained geometries similar to ref^11^. We remove molecules with consecutive double bonds, fused ring systems of more than four members or that form loops, and rings other than five- and six-membered rings. For consistency we apply this filtering to all compounds sets.

### Synthesizable Analog Search

We perform GED search using SmallWorld^22^, which is the default tool used for similarity search in the ZINC database^3^. SmallWorld uses pre-indexing of each library’s topological space as anonymous graphs which can then be used for rapid (sublinear) search of compounds with matching or similar topology. With the topological relationship between two compounds known, the GED calculation is greatly simplified. SmallWorld also calculates the Daylight fingerprint and ECFP4 fingerprint Tanimoto distances.

### Conformer Generation

Starting from the SMILES string representation of a molecule, we generate five conformers (3D coordinates) using the ETKDG algorithm in RDKit23. To reduce docking of redundant structures, we perform Butina clustering using RMSD as the distance metric on the five conformers, keeping only the centroid for each cluster. We then dock each remaining conformer to the corresponding target and keep the best scoring conformer as a representative.

### Ligand Docking

Docking compounds to target pockets is performed with QuickVina2^20^ (with default settings and a 20 Å wide bounding box), a computationally efficient version of AutoDock Vina^24^. Protein structures are prepared for docking using AutoDockTools^25^. Prior to docking, we perform force field optimization on all molecular structures using the MMFF94 force field^26^ in RDKit^23^. This step is especially important with de-novo generated structures, which are likely to have unrelaxed conformations. Using unrelaxed structures before docking could lead to significantly exaggerated docking scores.

### Evaluation

For evaluation, we chose 30 protein targets from the PDBbind general set. Our selection process includes: 1) clustering all protein sequences by 30% sequence identity, 2) randomly sampling 30 clusters and 3) choosing one random target per cluster. We generate 1000 molecules for each of the receptors in the test set using each of the three SBDD models. As a reference set for random virtual screening, we randomly sampled 50k compounds from the ZINC20 drug-like subset. We perform force field optimization on each of the molecules using MMFF94 in RDKit^23^ before virtually docking to the corresponding receptor. However, Vina score is strongly correlated with molecule size^11^ which prevents direct comparison between sets of compounds with different size distributions. For this reason, we primarily report Vina efficiency, or Vina score divided by number of heavy atoms, as it corrects for this size bias.

### Evaluation Properties and Metrics

Binding affinity is estimated for each ligand with the *Vina* docking score calculated with QuickVina2^20^, a speed optimized version of AutoDock Vina^24^. We define *ΔVina* as the difference between the Vina score of a generated ligand and a reference ligand (e.g., query ligand or ligand from crystal structure). *Vina* efficiency *score* (or ligand efficiency) is calculated as the *Vina* score divided by the number of heavy atoms. *Graph Edit Distance (GED)* is the number of edits needed to be made to the chemical structure of a compound to make it identical to a reference compound, and in this work is calculated using SmallWorld^22^. *Daylight distance* is calculated as one minus the *Tanimoto distance* between the Daylight fingerprints of a pair of molecules. *ECFP4 distance* is calculated as one minus the *Tanimoto distance* between the ECFP4 fingerprints of a pair of molecules. The *Quantitative Estimate of Drug-Likeness* (*QED*) score estimates the drug-likeness of a molecule by combining a set of molecular properties^28^. The *Synthetic Accessibility (SA)* score estimates the ease of synthesis of a molecule^29^.

## RESULTS

### Generated compounds tend to have synthesizable analogs

We start by generating approximately one thousand molecules for each of our test target pockets using three SBDD generative models: DrugHIVE^11^, DiffSBDD^13^, and Pocket2Mol^12^. We then dock each generated molecule into the corresponding target pocket using QuickVina2^20^. To select our final generated set, we filter out molecules with PAINS patterns, unusual structures that could cause inaccurate docking results (see Compound Filtering), or properties outside of the typical drug-like range^30^. We then rank each compound by normalized *Vina* score and take the 10 best compounds for each model and target. The properties of all generated molecules compared to the final set can be seen in Figure S1.

For each compound in the final generated query set, we perform a similarity search over existing commercially available and readily synthesizable compound libraries. For each generated query compound, we use the SmallWorld^22^ API to search for up to 1000 closest compounds by *GED* within a maximum *GED* of 12, for each of the Zinc^3^, Enamine REAL^2^, and Wuxi GalaXi^3^ libraries. We then rank the returned search compounds by *GED* and then *Daylight distance*, selecting the top-100 for our final search-hit set. We then generate 3D conformers and dock each final search-hit compound to the corresponding protein pocket.

The resulting top search-hit compounds are highly synthesizable and even tend to have improved predicted binding scores compared to the corresponding generated query compounds. As expected, we find a drastic improvement in predicted synthesizability of the top search-hit compounds when compared to the generated query compounds (Figure 2a). We also find an average improvement in predicted binding affinity compared to the query compounds (Figure 2b), with virtually all (98.7%) search-hit compounds within the Vina estimated margin of error of ±1.5 kcal/mol (Figure 2c). We also see that virtually all of the top search-hit compounds have equivalent or better predicted binding affinity compared to the corresponding PDB co-crystal ligand (Figure 2d), with a large number (38.8%) achieving significantly better scores (*ΔVina* < –1.5 kcal/mol). For all three models tested, we see a similar distribution shift toward improved predicted binding efficiency in the search-hit compounds, with an average improvement of the median Vina efficiency (Figure 2e).

**Figure 2.**
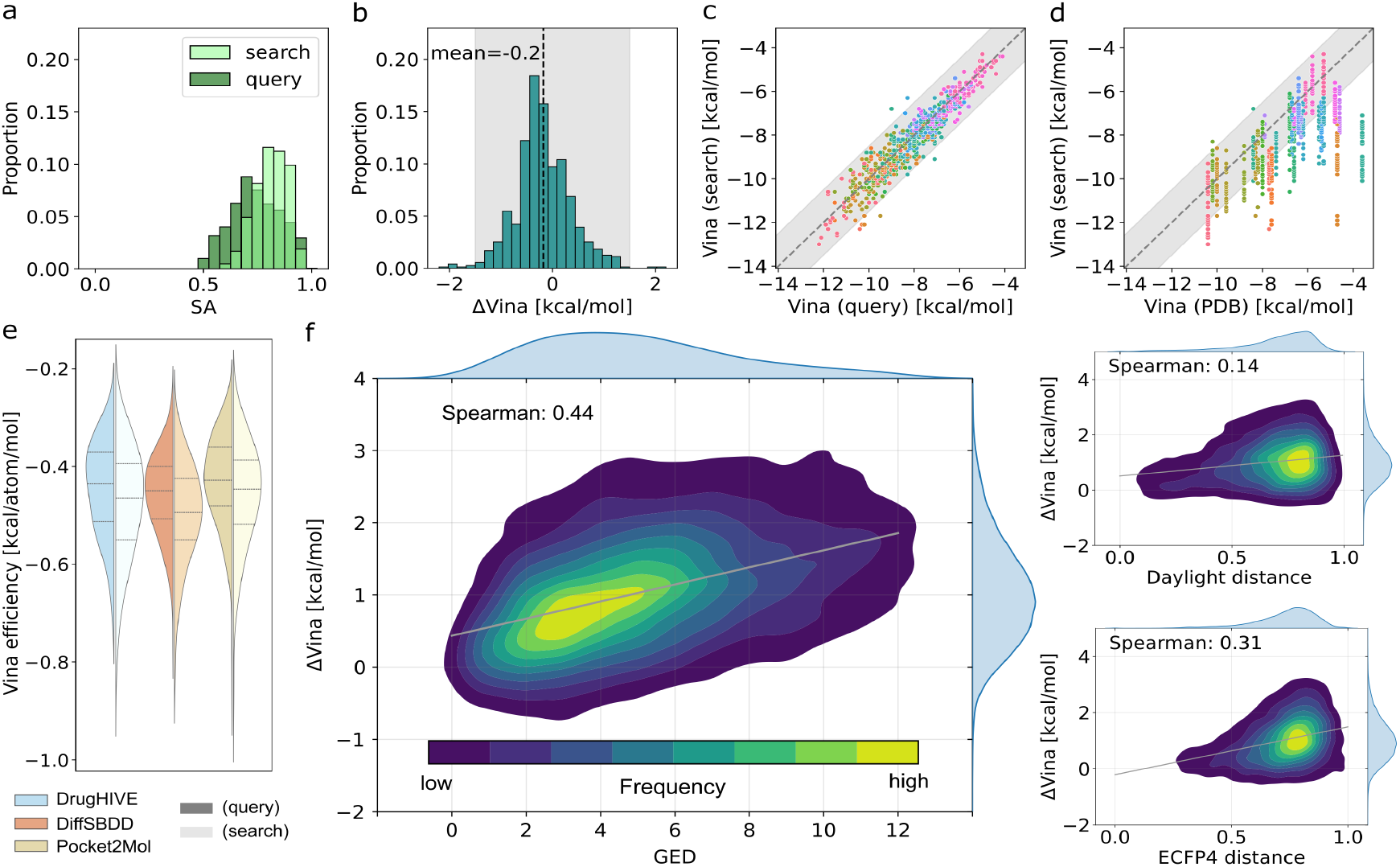
Generated compounds vs. search-hit compounds. (a) Histogram showing distribution of Synthetic Accessibility (SA) score for *generated query* and search-hit compounds. (b) Histogram of *ΔVina*^*\**^ scores for top-1 *search-hit* compounds with mean (black dashed line) and *Vina* uncertainty (shaded). (c) *Vina* scores for top-1 search-hit compounds versus the corresponding query compound for each target (colors) with *Vina* uncertainty (shaded). (d) *Vina* scores for top-1 search-hit compounds versus the corresponding crystal ligand for each target (colors) with *Vina* uncertainty (shaded). (e) Distributions of *Vina* efficiency for queries (dark) and top-1 search-hits (light) for each model (colors). (f) Distribution of *ΔVina* scorses for all (top-100) search-hits with respect to *graph edit distance* (*GED*), *Daylight distance*, and *ECFP4 distance*. Linear regression fit (grey line) shown along with computed Spearman (*ρ*) correlation coefficient. *\*ΔVina* = *Vina*_search-hit_ – *Vina*_query_

In Figure 2f, we show a positive Spearman correlation between *GED* and worse binding affinity scores (*ρ*=0.44). We observe similar, though weaker, correlations for *ECFP4 distance* (*ρ*=0.31) and *Daylight distance* (*ρ*=0.14). Since lower similarity search-hits from these databases tend to have worse binding scores than the query, this suggests that the generated query compounds indeed represent good templates for a successful predicted binder. Further, *GED* appears to be a better predictor of binding score than the two fingerprint-based similarity measures. This is likely due to GED being a better measure of topological and structural similarity, which are key determinants of shape complementarity and interaction geometry within the binding pocket.

### Generative SBDD paired with analog search outperforms random screening

To be practically useful, a generative approach combined with similarity search needs to outperform random virtual screening. To assess whether this is the case, we compare our generative approach results with screening of random compounds from the ZINC^31^ database. We randomly sampled 50k compounds from the Zinc drug-like compound subset and docked all compounds into each of our test pockets. In Figure 3a-b, we compare the Vina efficiency of the top-10 compounds from the generated queries, search-hits, and different size random Zinc subsets (ranging from 1k to 50k). The query compound distributions, which were selected from 1k generated compounds for each target, show significantly better Vina efficiency (Welch’s t-test *p*<0.05) than screening 10k (Pocket2Mol), 30k (DrugHIVE), and 50k (DiffSBDD) Zinc compounds. The search-hit compound distributions for all models show significant improvement (Welch’s t-test *p*<0.05) compared to their respective query compound distributions. Across all models, the top-10 similarity search-hits score better than those from screening up to 50k random Zinc compounds, despite only screening 2k compounds (including queries) per target—representing a 25x improvement in screening efficiency.

**Figure 3.**
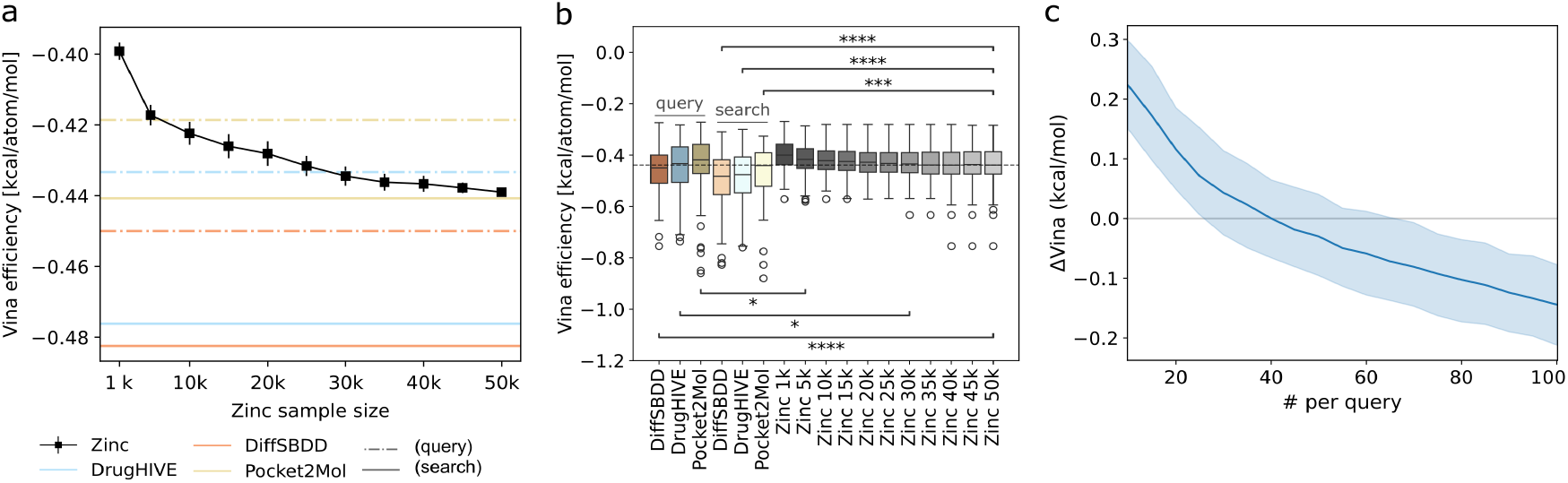
Comparison with random Zinc compounds. (a) Median Vina efficiency of the top-10 compounds for different size subsets of the random Zinc set relative to query (dashed) and search (solid) *hit* compounds for each model (color). Calculations for Zinc subsets are repeated a total of *n*=20 times, with median average and std. dev. shown. (b) Boxplots show the distributions of top-10 Vina efficiency for generated *query* compounds, search-hits, and Zinc (random) subsets of varying size. For each *query* or search-hit distribution, the largest Zinc subset distribution with significant Welch’s t-test result is indicated. Significance denoted as: *p* < 0.05 (∗*), p < 0*.*01 (*∗∗*), p < 0*.*001 (*∗∗∗), *p* < 0.0001 (∗∗∗∗). (c) Average *ΔVina* between top-10 scoring search-hit compound and *query* compound vs. number of total ranked search-hits that were docked to the target pocket, with 95% confidence interval (shaded).

These results represent selection from a docked candidate pool of the 100 most similar search-hits per query compound, however, docking and ranking fewer search-hits could lower the computational cost while still achieving favorable results. To assess this, we re-ran selection using different size search-hit candidate pools (docked search-hits) ranging between 1 and 100. In Figure 3c, we show the resulting *ΔVina* averaged across all models and targets ranges from -0.1 to 0.2, with parity (*ΔVina*=0) at approximately 40 search-hits docked per query. Surprisingly, even when using only a single search-hit, the upper limit of the *ΔVina* 95% confidence interval remains at a modest +0.3 kcal/mol—well within the estimated ±1.5 kcal/mol *Vina* uncertainty^24^. This demonstrates that synthesizable analogs with comparable docking scores can be reliably identified for generated compounds even with relatively few search-hits screened per query.

### Shared intermolecular interactions between predicted binding poses

Our results so far have shown that using similarity search to identify synthesizable analogs to *de novo* generated compounds reliably results in highly similar compounds with strong predicted binding affinity. However, this does not guarantee that the search-hit compounds have a similar predicted binding pose within the target pocket. To assess this, we compared the docked intermolecular interactions made by the top predicted binding pose of the search-hit compounds to those made by corresponding generated query compound. Interactions were identified using the Protein-Ligand Interaction Profiler (PLIP)^27^ on the top ranking docked pose from QuickVina2^20^ for each compound. We consider matching interactions between query compound and search-hit compound at two levels of specificity: 1) same target residue and 2) same specific target atom. In Table 2, we show a summary of the proportion of search-hits having shared interactions with the generated query compound for each interaction type (H-bond, hydrophobic, salt bridge, π-cation, and π-stacking) across all models and targets. We observe that a majority of search-hits share at least one interaction with the generated query across all interaction types. We also see that search-hits are less likely to share the more frequent H-bond and hydrophobic interactions than the less frequent ones, except for π-stacking which is both frequent and highly conserved.

**Table 1:** Protein-ligand interaction cutoff criteria. For each interaction type, we list the PLIP^27^ non-default cutoff criteria that we use for identifying intermolecular interactions. All table values are distance cutoffs, except for hydrogen bonds. For hydrogen bonds, we choose a stricter distance and D-H-A angle cutoff. In the case of π interactions, the table values refer to center–center or center–cation distance.

| H-bond | Hydrophobic | Salt bridge | π-stack (parallel) | π-stack (perpendicular) | π-cation |
| --- | --- | --- | --- | --- | --- |
| 3.5 Å, 120° | 3.8 Å | 4.0 Å | 4.0 Å | 5.5 Å | 4.0 Å |

**Table 2:** Shared intermolecular interactions by interaction type. Proportion of search-hits having shared interactions with the generated query compound across all models and targets. Statistics are reported for search-hits that share all or at least 1 of the query interactions, as well as for different positive match types: matching interactions share the same 1) target atom or 2) target residue. For each interaction type the total number (*n*) of search-hit compounds with a query compound predicted to make the corresponding interaction with the target is reported.

| #<br>matched | Match<br>type | H-bond<br>(*n*=36404) | Hydrophobic<br>(*n*=62582) | Salt bridge<br>(*n*=1332) | π-cation<br>(*n*=474) | π-stacking<br>(*n*=29830) |
| --- | --- | --- | --- | --- | --- | --- |
| *all* | atom | 0.17 | 0.12 | 0.58 | 0.79 | 0.57 |
|  | residue | 0.27 | 0.30 | 0.58 | 0.79 | 0.83 |
| *at least 1* | atom | 0.65 | 0.75 | 0.73 | 0.79 | 0.90 |
|  | residue | 0.69 | 0.86 | 0.73 | 0.79 | 0.92 |

In Figure 4, we show the proportion of docked search-hit compounds that share at least one or all specific query compound interactions for each generative model. Here we choose to focus on specific non-hydrophobic interactions (Hydrogen bonds, π interactions, and salt bridges), because we are primarily interested in assessing how often search-hits exhibit the same type of target specificity as the query. The same statistics for all interactions can be found in Figure S2. In Figure S3, we show the distribution of intermolecular interactions made by all generated query compounds. The average number of intermolecular interactions made by query compounds is 2.5 (specific) and 5.2 (all). Across all query compounds, models and targets, an average of 19.5% (atom match) to 31.4% (residue match) of search-hits (top-100) share all specific interactions with the query and 76.7% (atom match) to 79.5% (residue match) share at least one (Figure 2a). For top-1 search-hits (Figure 2b), we see that a majority of queries have at least one search-hit sharing all specific interactions (50.9% and 80.7% for atom and residue match, respectively) and nearly all queries have at least one search-hit that shares an interaction (99.2% for both atom and residue match). In Figure 4c, we show that there is some variation in the proportion of queries with search-hit compounds matching all interactions across the different protein targets. However, even for the lowest performing target (PDB ID 5G1P) 10.5% (atom match) to 20.0% (residue match) of query compounds have an exact interaction analog. For all targets, nearly every query compound has a search-hit with at least one interaction.

**Figure 4.**
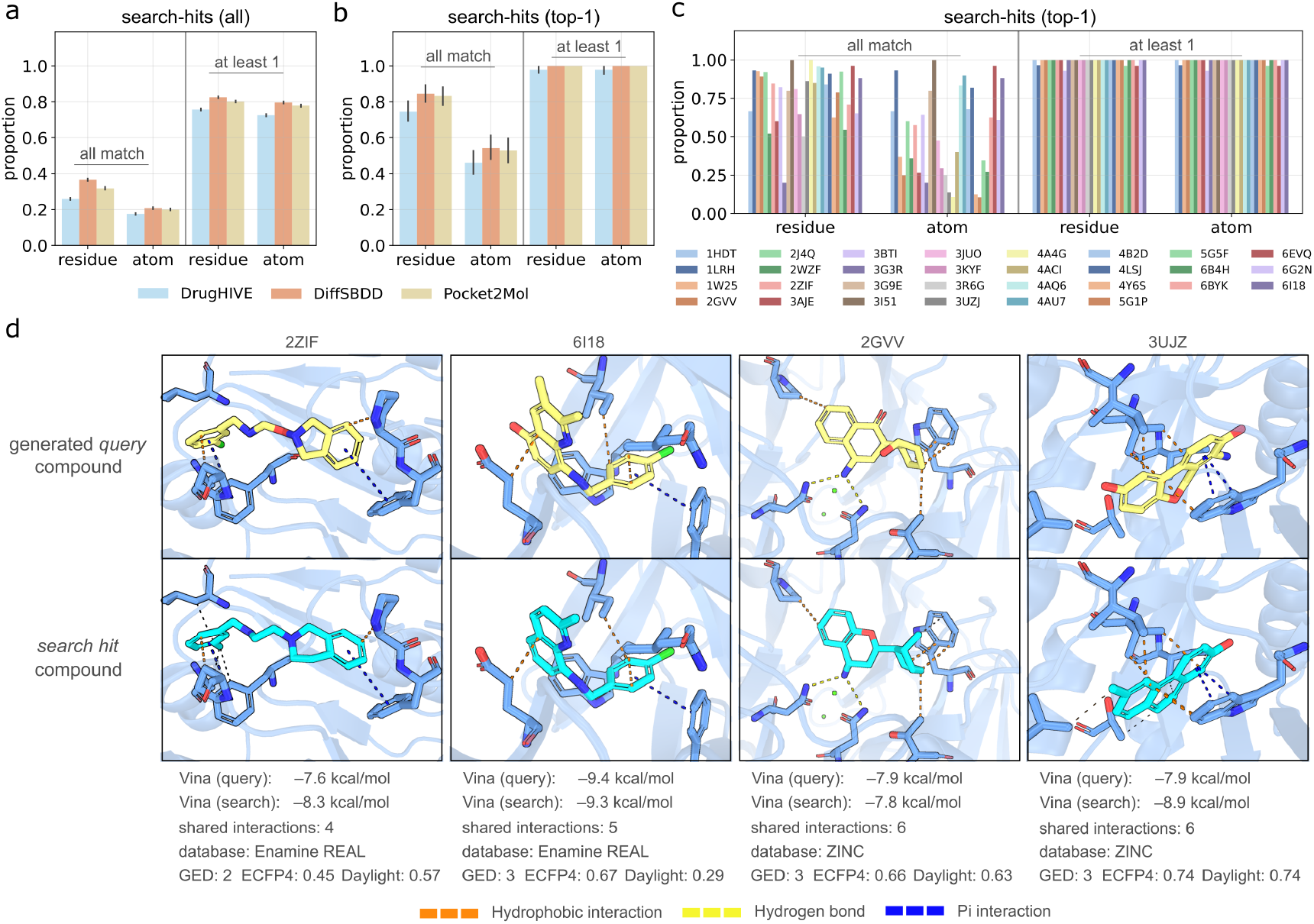
Shared intermolecular interactions between generated queries and search-hits. (a-c) Bar plots showing proportion of search-hits that share *all* or *at least 1* of query specific protein–ligand interactions. Proportions are shown for both exact residue match (residue) and exact atom match (atom) criteria. (a) Proportion of top-100 *search-hits* for each query with matching specific interactions for each model. (b) Proportion of top-1 *search-hits* for each query with matching specific interactions for each model. (c) Proportion of top-1 *search-hits* for each query with matching specific interactions by protein target across all models. (d) Docked poses of *search-hit* (yellow carbon) and *query* (cyan carbon) compound pairs with protein-ligand interactions shown. Vina docking score, number of shared interactions, search-hit database, and chemical distance metrics (*GED, ECFP4 distance, Daylight distance*) are shown for each pair.

To give a better sense of what the docked poses of interaction analogs look like, in Figure 4d we show a selection of docked poses with highlighted intermolecular interactions for query and search-hit pairs with all interactions shared (atom match) for four of the targets. We report the *Vina* scores of each compound, the number of shared interactions, database of the search-hit, and the molecular distance metrics. We observe that search-hits with many shared query contacts tend to also have similar predicted binding poses and a relatively low *GED*. Interestingly, fingerprint-based distance metrics do not always agree with *GED* (or with each other). For example, the pair of compounds displayed for target PDB ID 3JUZ have clear structural similarities and a relatively low *GED* (3) but high *Daylight distance* and *ECFP4 distance* (0.74 for both). For the pair of compounds displayed for target PDB ID 6I18, *GED* (3) and *Daylight distance* (0.29) are low while *ECFP4 distance* (0.67) is high. To quantify this, we calculated the Spearman correlation between each of the molecular distance metrics and the number of shared interactions (atom match). We found that *GED* has the highest correlation, especially as the number of protein–ligand interactions made by the query compound increases (Figure S4). These results are consistent with the earlier findings (Figure 2f) that *GED* is a better predictor of binding affinity score than the fingerprint-based similarity metrics.

## DISCUSSION

In this work, we have shown that current generative structure-based drug design (SBDD) models paired with similarity-based search can be used to discover synthesizable compounds with high predicted binding affinity for a wide range of protein targets. Generative models tend to produce compounds with low synthetic accessibility scores^14–16^, which is a key limitation to practical drug discovery^4^. To address this, we show that model-guided virtual screening (MGVS)—using generated compounds to guide the search for synthesizable analogs—effectively overcomes this limitation. We apply MGVS across 30 diverse protein targets and observe that for three separate SBDD models (DiffSBDD^13^, DrugHIVE^11^, and Pocket2Mol^12^) that screening only 2k compounds leads to better quality candidates than standard VLS screening of 50k ZINC compounds, representing a 25x improvement in efficiency.

Although current generative models tend to produce compounds with low synthesizability, they are effective at identifying compounds located in promising regions of chemical space. Once these regions have been identified, similarity search can be used to find adjacent synthesizable analogs. Across all models and targets, we find that virtually all (98.7%) generated compounds selected for analog search yielded a search-hit with equivalent or better docking scores. We also found that search-hits commonly have similar docking poses to the query compound, indicating that the compounds produced by generative models help identify promising potential binding modes. We observe that ∼50% of query compounds yielded a search-hit with a docking pose sharing all specific (non-hydrophobic) residue-level protein-ligand interactions, and virtually all (∼99%) yielded a search-hit sharing at least one interaction (Figure 4b). Assuming that the number of shared intermolecular interactions is an indicator of docking pose similarity (Figure 4d), this demonstrates consistent success in identifying predicted binding analogs.

Altogether these results demonstrate that current generative models, despite their limitations, can already be used to improve virtual screening efficiency through MGVS. We show that identifying compounds with strong predicted affinity for a protein target, regardless of synthesizability, is useful for guiding the screening of high-potential subsets of existing ultra-large chemical databases. To be successful, this approach relies on search methods efficient enough to operate on these large and growing chemical spaces^1^, which could be viewed as a limitation. However, improving chemical search methods is an active area of research^32,33^ that should enable efficient querying of expanded chemical databases. As chemical databases grow and search methods improve, we would expect MGVS to become even more effective due to improved retrieval. On the contrary, the expansion of chemical spaces poses a real challenge to existing VLS methods that rely on exhaustive screening^9^. Even for the most efficient synthon-based VLS approaches used to screen current ultra-large libraries, scaling to chemical spaces orders of magnitude larger will require innovations that surpass what current methods can support.

There are efforts to overcome the generative SBDD synthesizability issue by designing models restricted to synthesizable subspaces^17–19^. However, we observe that even for the current (unrestricted) models that we tested, while they are able to generate high quality binding candidates, the set of synthesizable analogs identified based on these initial queries often contain compounds with even better predicted binding. This suggests that current models cannot consistently generate the most optimal compounds within these localized regions of chemical space and restricting them further could lead to even worse performance. Efforts to improve the ability of models to generate ideal binders, regardless of synthesizability, seem to be warranted. Applying current models in MGVS already achieves a ∼25x improvement in screening efficiency over standard virtual screening, so further improvement to models that can be utilized in this paradigm could have significant impact on early drug discovery.

## ASSOCIATED CONTENT

### Supporting Information

Property distributions of generated compounds; shared intermolecular interactions between search queries and hits for all interaction types (including hydrophobic); distributions of number of protein-ligand interactions for docked poses across all generated query compounds; correlation of molecular similarity metrics with number of shared interactions between query and search-hit compounds (PDF)

## AUTHOR INFORMATION

### Authors

**Jesse A. Weller -** Department of Quantitative and Computational Biology, Department of Physics & Astronomy, University of Southern California, Los Angeles, CA 90089, USA;

**Jinsen Li -** Department of Quantitative and Computational Biology, University of Southern California, Los Angeles, CA 90089, USA;

**Yibei Jiang -** Department of Quantitative and Computational Biology, University of Southern California, Los Angeles, CA 90089, USA;

### Author Contributions

Project conception (J.A.W., R.R.), lead project design (J.A.W.), supporting project design (J.L., Y.J., R.R.), data generation and analysis (J.A.W.), manuscript writing (J.A.W.), manuscript edits (J.A.W., J.L., Y.J., R.R.), project supervision and funding (R.R.).

### Funding Sources

This work was supported by the National Institutes of Health [grant R35GM130376 to R.R.] and a University of Southern California Office of Research and Innovation SBIR/STTR Planning Award [to R.R.].

## ACKNOWLEDGEMENT

The authors acknowledge helpful discussions with other members of the Rohs lab.

## ABBREVIATIONS

ALogP: octanol-water partition coefficient (Crippen)
HBA: hydrogen-bond acceptors
HBD: hydrogen-bond donors
HTS: high-throughput screening
MGVS: model-guided virtual screening
PAINS: Pan-Assay Interference Compounds
PDB: Protein Data Bank
QED: Quantitative Estimate of Drug-likeness
RMSD: root mean square distance
SA: Synthetic Accessibility
SAR: structure-activity relationship
SBDD: structure-based drug design
SMILES: Simplified Molecular Input Line Entry System
TPSA: topological polar surface area
VAE: variational autoencoder
VLS: virtual ligand screening;

## DATA AND SOFTWARE AVAILABILITY

Data used in evaluation including molecules, molecular properties, and docking scores, are available in supplementary information.

## Notes

### Competing Interest Statement

The authors have declared no competing interest.

