## Supplemental Information for "Circumventing the synthesizability problem in generative molecular design"

Supplementary Figures S1–S4.

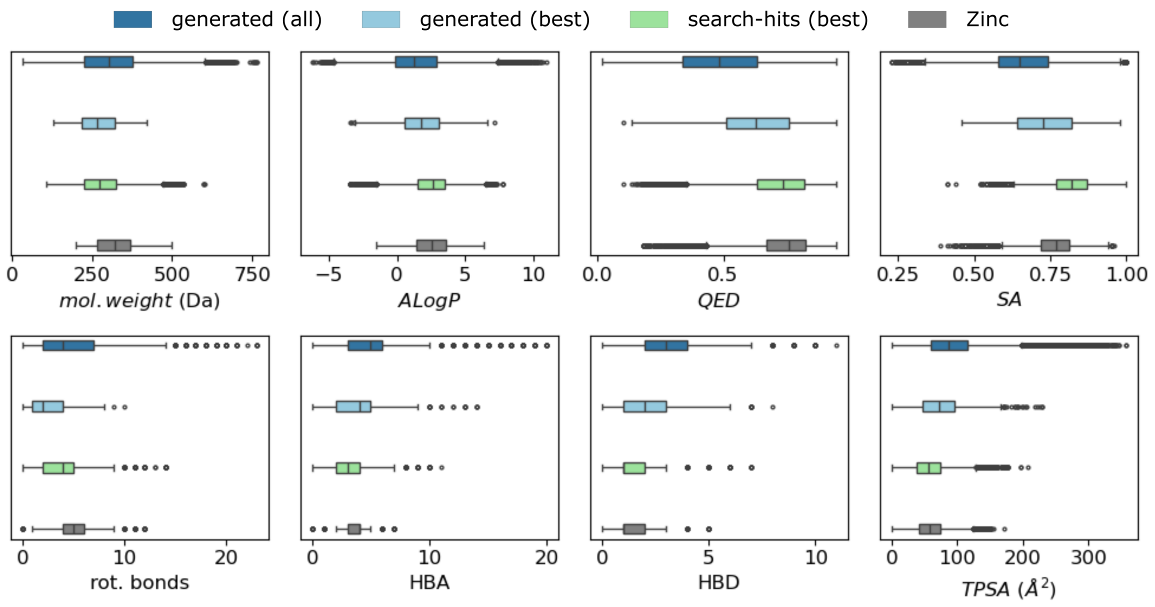


**Figure S1.** Property distributions of generated compounds. Comparison of all generated compounds, best generated (query) compounds, search-hit compounds, and random Zinc compounds. (mol. wt.: molecular weight, SA: synthetic accessibility, QED: drug-likeness, rot. bonds: rotatable bonds, HBD: Hydrogen bond donors, HBA: Hydrogen bond acceptors, TPSA: topological polar surface area, alogP: hydrophobicity).

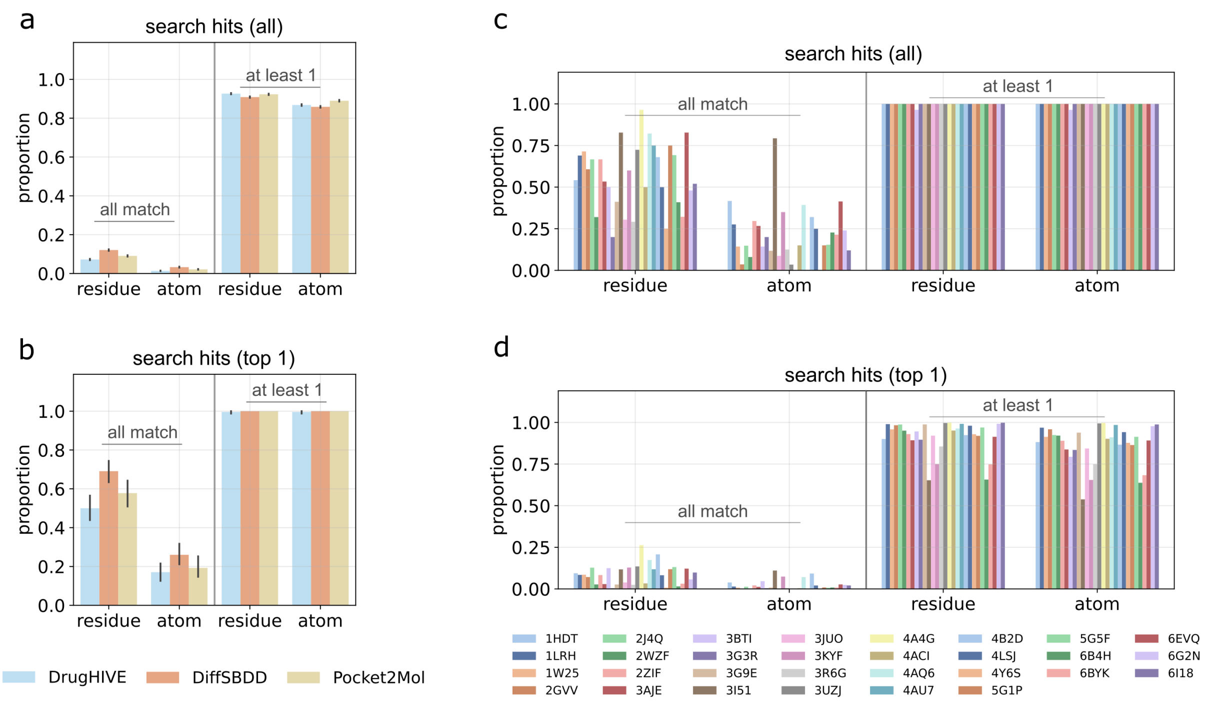


**Figure S2.** Shared intermolecular interactions between search queries and hits for all interaction types (including hydrophobic). (a-b) Bar plots showing proportion of search hits that share *all* or *at least 1* of query protein–ligand interactions. Proportions are shown for both exact residue match (residue) and exact atom match (atom) criteria. (a) Proportion of top-100 *search hits* for each query with matching interactions for each model. (b) Proportion of top-1 *search hits* for each query with matching interactions for each model. (c) Proportion of top-100 *search hits* for each query with matching interactions by protein target. (d) Proportion of top-1 *search hits* for each query with matching interactions by protein target.

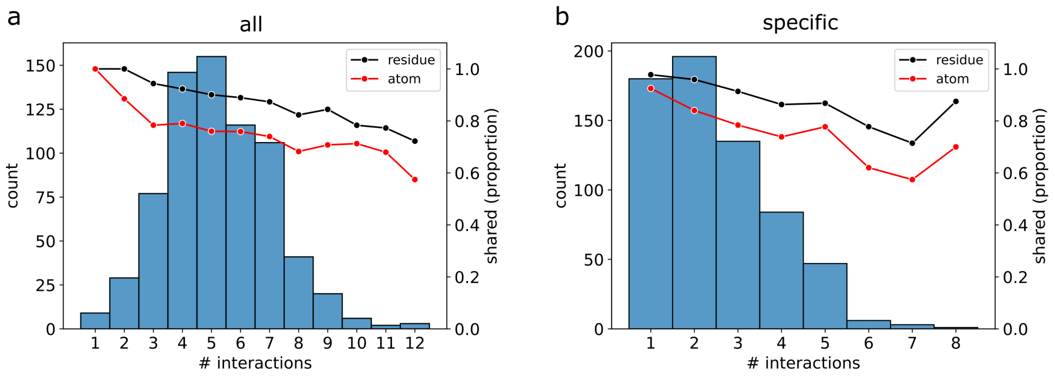


**Figure S3.** Distributions of number of protein–ligand interactions for docked poses across all generated query compounds. Histograms show the interaction counts and lines show average proportion of shared interactions (residue-match, red; atom-match, black) of the top search hit for each query. Separate plots for (a) all interactions (including hydrophobic) and (b) specific interactions only (excluding hydrophobic) are shown.

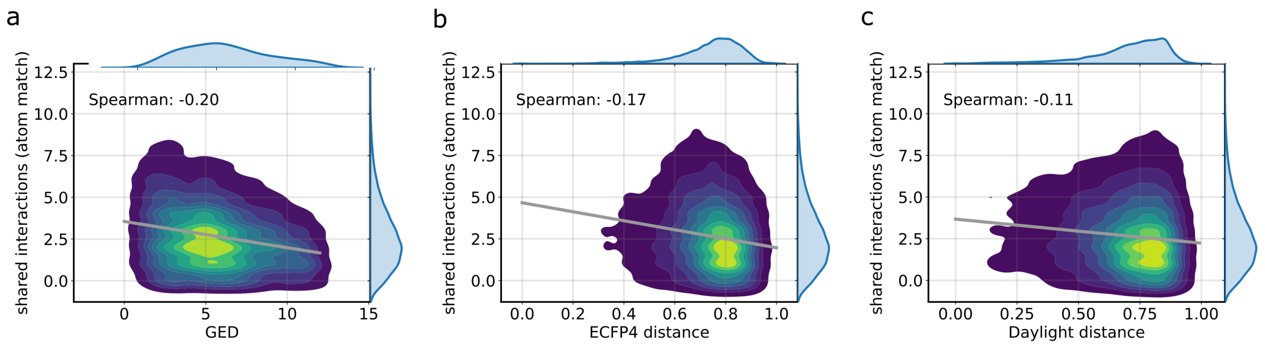


**Figure S4.** Correlation of molecular similarity metrics with number of shared interactions between query and search-hit compounds for (a) *graph edit distance* (*GED*) (b) *ECFP4 fingerprint Tanimoto distance* and (c) *Daylight fingerprint Tanimoto distance*.
